# The Mixed Blessing of AMPK Signaling in Cancer Treatments

**DOI:** 10.1101/2021.06.25.449964

**Authors:** Mehrshad Sadria, Deokhwa Seo, Anita T. Layton

## Abstract

Nutrient acquisition and metabolism pathways are altered in cancer cells to meet bioenergetic and biosynthetic demands. A major regulator of cellular metabolism and energy homeostasis, in normal and cancer cells, is AMP-activated protein kinase (AMPK). AMPK influences cell growth via its modulation of the mechanistic target of Rapamycin (mTOR) pathway, specifically, by inhibiting mTOR complex mTORC1, which facilitates cell proliferation, and by activating mTORC2 and cell survival. Given its conflicting roles, the effects of AMPK activation in cancer can be counter-intuitive. Prior to the establishment of cancer, AMPK acts as a tumor suppressor. However, following the onset of cancer, AMPK has been shown to either suppress or promote cancer, depending on cell type or state. To unravel the controversial roles of AMPK in cancer, we developed a computational model to simulate the effects of pharmacological maneuvers that target key metabolic signalling nodes, with specific focus on AMPK, mTORC, and their modulators. Model simulations clarify the competing effects and the roles of key metabolic signalling pathways in tumorigenesis, which may yield insights on innovative therapeutic strategies.

## Introduction

Controlled cell division is essential for cellular homeostasis and normal development. Abnormal cell proliferation is associated with pathological states such as cancer. To support their growth and proliferation, cancer cells reprogram their metabolism to promote nutrient uptake (1). Despite the heterogeneity in genetic mutations that leads to tumorigenesis, a common alteration in tumors occurs in pathways to upregulate nutrient acquisition and optimize nutrient utilization when resources are scarce (2). Indeed, metabolic reprogramming is a hallmark of cancer, and a better understanding of the synergy among the many signalling proteins and kinases may yield useful targets for therapies.

One of the central signaling pathways that control metabolic processes is the mechanistic target of Rapamycin (mTOR) pathway. mTOR is a highly conserved serine/threonine protein kinase with two distinct complexes, mTORC1 and mTORC2. mTOR controls cell growth, proliferation, motility and survival, protein and lipid synthesis, glucose metabolism, mitochondrial function and transcription, in response to nutrient and hormonal signals (3). mTOR is estimated to be hyperactivated in over 70% of cancers (4) and its hyperactivation leads to tumor growth metastasis and angiogenesis (5). This has stimulated interest in targeting mTORC1 for cancer therapy. Notably, rapamycin and its analogs bind to a domain separate from the catalytic site to block a subset of mTOR functions, inhibiting cell growth (6). A limitation of rapamycin is that it is only an allosteric mTOR inhibitor and does not fully block its activity. Second generation mTOR inhibitors that target the catalytic site of mTOR to block mTOR activity have been developed (7). However, while the potency of these inhibitors in slowing cell growth and proliferation has been demonstrated, their toxicity has limited their usage (8). Furthermore, the metabolic plasticity of cancer cells allows them to trigger mTOR-independent mechanisms to compensate for the inhibited mTOR activity, thereby allowing the cells to acquire sufficient nutrients for growth and proliferation. Consequently, there is an urgent need to improve therapeutic strategies to maximize their benefits (9).

Among the several proteins associated with mTOR and the formation of mTORC1 and mTORC2, one particularly intriguing modulator is the DEP domain-containing mTOR-interacting protein (DEPTOR). Taken in isolation, DEPTOR inhibits both mTORC1 and mTORC2 (10). However, when feedback loops are taken into account, the overall impact of DEPTOR on mTOR signaling becomes more complex (11). While DEPTOR depletion promotes mTORC1 activity, its loss has been reported to unexpectedly inhibit mTORC2 (10). Conversely, overexpression of DEPTOR reduces mTORC1 signaling and promotes mTORC2 activity (10). This apparent paradox can be attributed to the mTORC1/Phosphoinositide 3-kinases (PI3K) feedback loop, whose opposing effect on mTORC2 dominates over DEPTOR. Because DEPTOR affects the signalling of mTOR and PI3K, which are key modulators of cell growth, survival, and proliferation, the relationship between DEPTOR and cancer is a promising research focus. But as is often the case in cancer research, the ideal target compounds may differ depending on cancer types. In most cancers, DEPTOR expression is low (12–16). Thus, it may be beneficial to administer compounds that enhance the stability of DEPTOR or its connection to mTOR, in an attempt to suppress mTOR and the growth, survival, and proliferation of cancer cells. However, in some other cancers, e.g., multiple myeloma, DEPTOR expression is elevated (10). Here, compounds that impair the DEPTOR-mTOR connections may be beneficial.

Another major regulator of cellular metabolism and energy homeostasis, in both normal and cancer cells, is AMP-activated protein kinase (AMPK). Known as a master regulator of cellular energy, AMPK is a heterotrimeric kinase complex consisting of a catalytic α-subunit and two regulatory subunits, β and γ (17). Under conditions of low energy, AMPK phosphorylates specific enzymes and growth control nodes to increase ATP generation and decrease ATP consumption. The link between AMPK and cancer was first identified through the tumor-suppressive function of liver kinase B1 (LKB1), which phosphorylates AMPK and leads to mTORC1 inhibition (18). Studies in animal models and humans have suggested that compounds that activate AMPK have health-promoting effects, including improvements in diabetes, cardiovascular health, and mitochondrial disease, and even extension in life span (19–22). However, the role of AMPK in tumorigenesis is controversial: while studies have suggested that AMPK suppresses cancer cell proliferation and tumor formation (23), recent evidence also suggests that AMPK can promote T-ALL tumor cell growth through cell energy maintenance in some model organisms (24,25). Further complicating the picture, a direct link between AMPK and mTORC2 was recently discovered, which reveals the ability of AMPK to promote cell survival under energetic stress situations and emphasizes the strong interplay between anabolism and catabolism regulators: AMPK and the mTORC family (26).

Given its multifaceted and controversial roles in cancer and its beneficial capacity in many diseases such as diabetes, under what conditions is it advantageous to activate AMPK and when is it not? How might a DEPTOR inhibitor differentially modify a cancer cell’s propensity to proliferate and persist? And how are these effects impacted by the cell’s microenvironment? The multitude of biochemical reactions and feedback loops involved in the metabolic pathway are difficult to untangle. As such, we developed a computational model to simulate the effects of pharmacological maneuvers that target key metabolic signalling nodes, with specific focus on AMPK, DEPTOR, and mTORC. Model simulations clarify some of these conflicting effects and provide a better understanding of the role of key metabolic signalling pathways in tumorigenesis. These findings may yield insights on innovative therapeutic strategies, including immunotherapy that targets relevant metabolic networks in cancer.

## Results

### Model Calibration and Validation

Model parameters are chosen so that the model predicts oscillatory time-profiles of mTORC1, ULK1 and AMPK consistent with experimental observations under energetic stress (27) (see materials and methods). The predicted time-courses are shown in Fig. 1(A-C) together with experimental data (27). The period of oscillation is approximately 24 hours. Additionally, model parameters are chosen to yield mTORC1 response to changes in AMPK activation level that match experimental values (28). The predicted relation is shown in Fig. 1D. Recently, it has been shown that AMPK promotes mTORC2 (26). Additional calibration is performed using AICAR experimental data. Upon injection, AICAR activates AMPK, which results in the phosphorylation of mTORC2, reflecting in AKT activation. A comparison between model predictions and experimental data (26,29) can be found in Fig. 1E.

**Figure 1.**
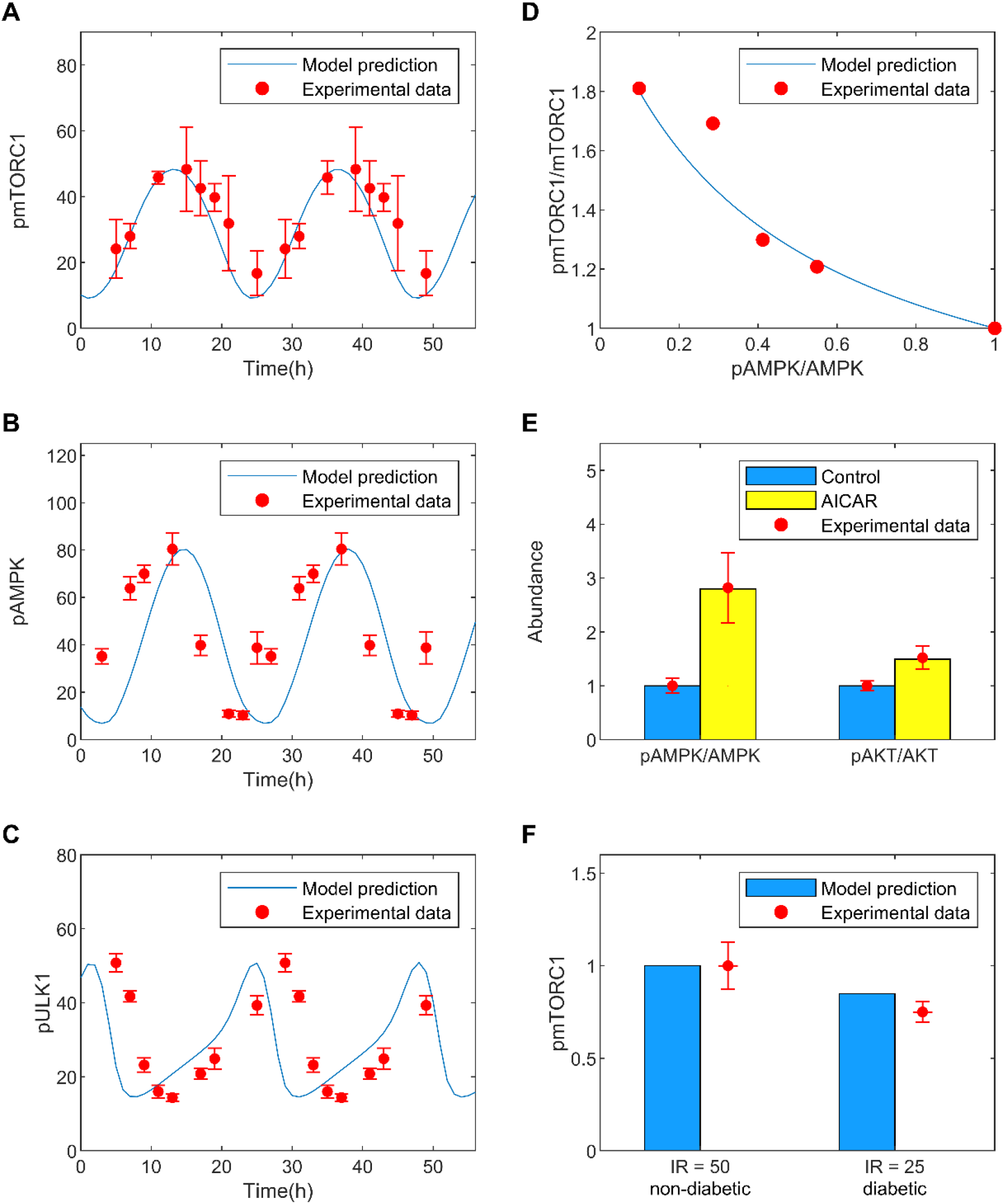
Model parameters are fitted against experimental data. *A-C*, oscillatory model solution under energetic stress. Predicted time profiles are shown for pmTORC1, pAMPK, and pULK1, together with the corresponding experimental data points (27). *D*, predicted phosphorylated-to-unphosphorylated mTORC1 ratio as a function of AMPK activation, compared with data from (28). *E*, predicted activation of AMPK and AKT following AICAR administration, compared with data from (26,29). *F*, predicted abundance of phosphorylated mTORC1 at different insulin receptor levels, corresponding to healthy and diabetic conditions. Diabetes values are normalized by the respective non-diabetes values. Experimental data (30) are shown in red dots with error bars.

To validate the calibrated model, simulation results were then compared against experimental data not used in the parameter fitting. An additional validation test was done using the observed reduction in mTORC1 activation in patients with diabetes. A diabetes model was simulated by decreasing total insulin receptor abundance by 50%. The model predicted fractional reduction in activated mTORC1 in diabetes that is consistent with clinical data (30); see Fig. 1F.

### Sensitivity analysis

Local sensitivity analysis can be used to understand the sensitivity of model outputs to local variations in individual parameters around baseline values. Most parameters have negligible local effect on key model outputs, with sensitivity values close to zero (Fig. 1S, supplemental materials). The lone exception is K_pmTORC1, which denotes the deactivation rate of mTORC1. Most outputs are highly sensitive to changes in K_pmTORC1, including IRS, AKT, AMPK, ULK1, SIRT1, mTORC1 and mTORC2. The sensitivity of model outputs to variations in K_pmTORC1 provides evidence for the key role of mTORC1 in determining network dynamics.

Global sensitivity analysis assesses the robustness of the model output to uncertainty in individual parameters over their entire range of interest. Results of the analysis are illustrated as a heatmap in Fig. 2A. The probability density functions of the sampled outputs are included in Fig. 2S. Results of this analysis can identify the most effective parameters to target to manipulate the abundance of a given protein or sets of proteins. Specifically, we seek to identify drug targets for suppressing the proliferation and survival of cancer cells. Is there a parameter to which mTORC1 and mTORC2 are highly sensitive, in the same direction? Such a parameter would indicate an ideal cancer drug target. Unfortunately, the global sensitivity results do not reveal such a parameter (Fig. 2A). As an alternative, we seek parameters that mTORC2 is highly sensitive to, as these can serve as drug targets to be administered in conjunction with an mTORC1 inhibitor. Parameters with opposite signs for the PRCC values of mTORC1 and mTORC2 were excluded, to avoid promoting mTORC1. The analysis indicated V_mTORC2 (deactivation rate of mTORC2) and Kd_mTORC2_DEPTOR (dissociation rate of the mTORC2-DEPTOR complex) as the most promising candidates for mTORC2 suppression.

**Figure 2.**
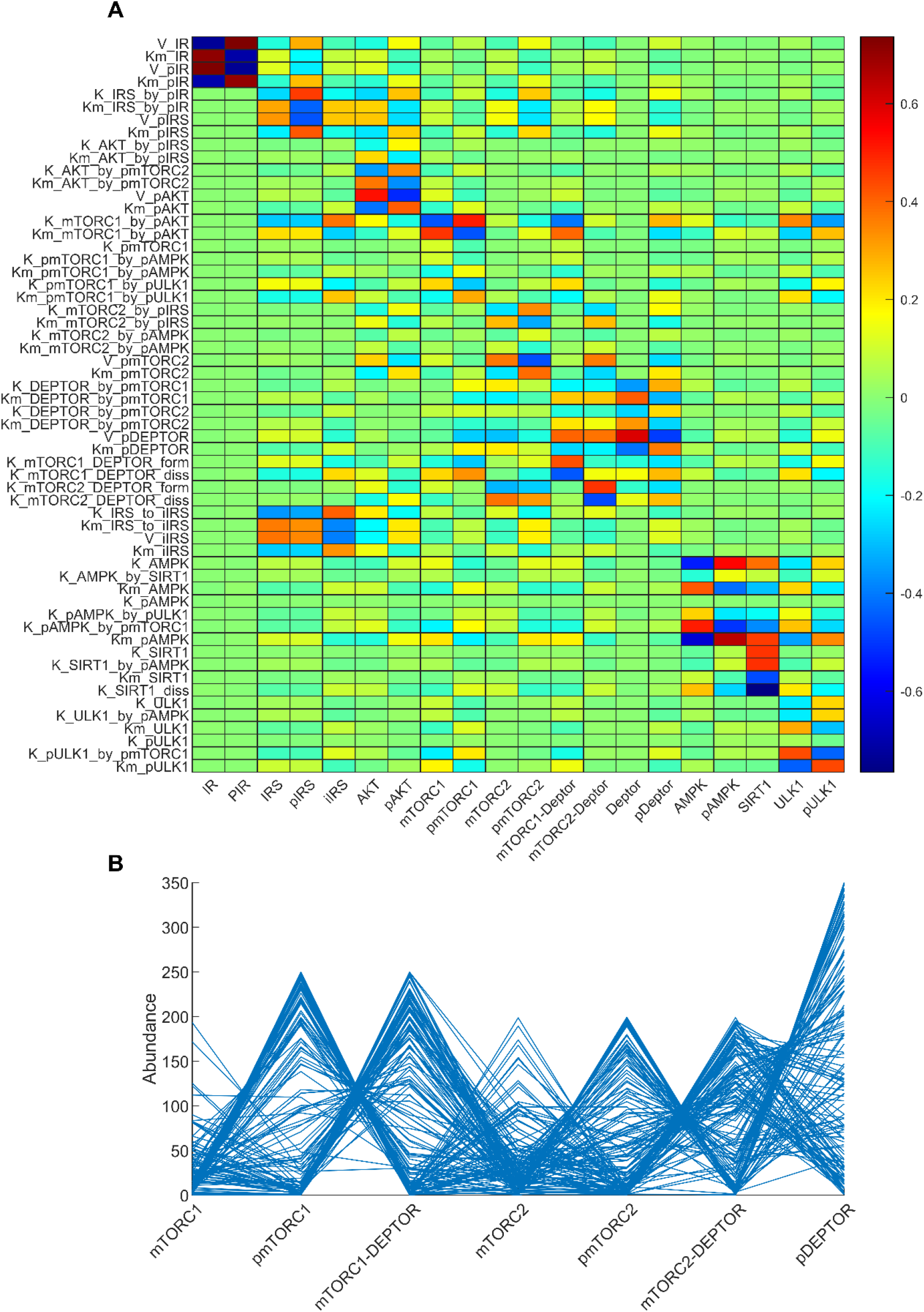
Top panel, heat map that illustrates the global sensitivity of key model outputs (horizontal axis) to variations in selected model parameters (vertical axis). Definition of the parameters can be found in the Supplemental Materials. Bottom panel, parallel coordinate plot showing the oscillations-inducing sets returned from a 7D analysis where the abundances of the model species are randomly sampled.

Furthermore, parameters that contribute the most to the variations in key model outputs were identified. mTORC1 is most sensitive to K_mTORC1_by_pAKT and Km_mTORC1_by_pAKT, in opposite directions, with PRCC values of −0.462 and 0.4671 respectively. These parameters constitute the Michaelis Menten kinetics that characterize the activation of mTORC1 by AKT (see model equations in Table 1S). Due to mTORC1’s strong inhibition of ULK1, ULK1 is similarly sensitive to variations in these parameters (Fig. 2A). Additionally, ULK1 is sensitive to variations in K_pULK1_by_pmTORC1, which characterizes the inhibition of ULK1 by mTORC1, and Km_ULK1, the Michaelis-Menton constant that characterizes ULK1 activation; the associated PRCC values are 0.4347 and −0.4385, respectively. AMPK is most sensitive to K_pAMPK_by_pmTORC1, which characterizes the inhibition of AMPK by mTORC1, and Km_AMPK, the Michaelis-Menton constant that characterizes AMPK activation; the associated PRCC values are 0.4993 and −0.6441, respectively. The former indicates the strong negative feedback strength of mTORC1 on AMPK.

To illustrate how the abundances of model proteins change concomitantly as model parameters vary, we randomly sampled 150 solutions and plotted the values of 7 key proteins in Fig. 2B (mTORC1, pmTORC1, pmTORC1+DEPTOR, mTORC2, pmTORC2, pmTORC2+DEPTOR, and pDEPTOR). Since total mTORC1 is conserved, a high phosphorylated mTORC1 level is associated with lower mTORC1 and mTORC1-DEPTOR. A similar relationship can be seen among the mTORC2 species and among the DEPTOR species.

### Is AMPK a cancer suppressor or promoter? It depends on cellular nutrient level

Activation of AMPK is generally viewed as beneficial. However, given AMPK’s impact on cellular energy and cycle, as well as the heterogeneity of cancer cell types, are there microenvironments under which AMPK activation may be harmful to the organism’s overall survival? To investigate this possibility, we simulated cellular microenvironments with different metabolic stress levels by considering a range of V_IR values. V_IR determines the activation level of IR; consequently, higher V_IR values correspond to higher insulin and thus glucose levels in the microenvironment, and vice versa. For each V_IR value we varied AMPK abundance, and analyzed changes in key variables mTORC1 and mTORC2 to assess the effects on cell population. Results are shown as a surface plot in Fig. 3B. We are particularly interested in the implications in cancer. Because mTORC1 and mTORC2 promote cell proliferation and survival, respectively, the simultaneous inhibition of both proteins may help to limit the proliferation of cancer cells.

**Figure 3.**
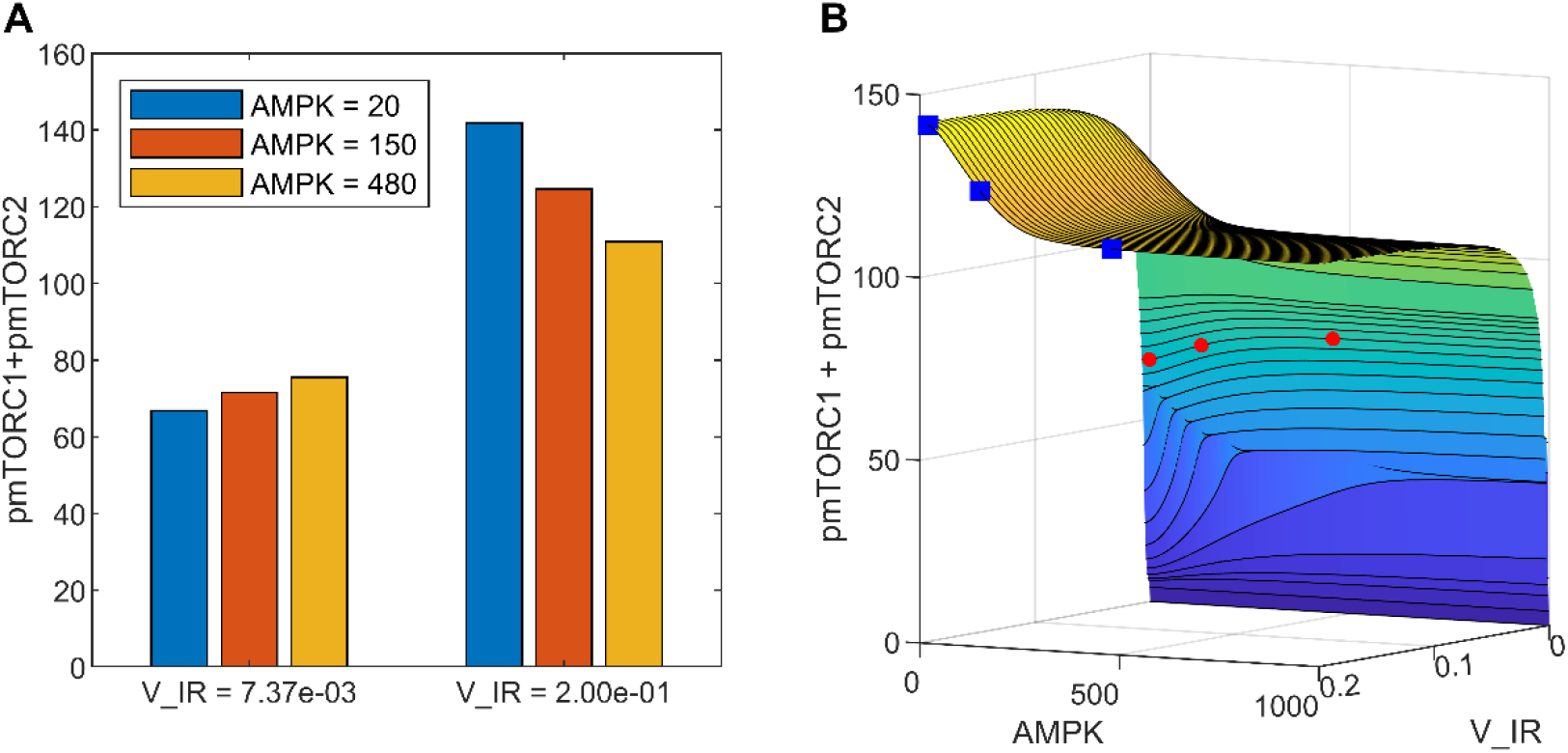
The effect of AMPK activation and nutrient levels on cell proliferation and survival, given by the sum of phosphorylated mTORC1 and mTORC2. *A*, results shown for two nutritional (V_IR) levels and three AMPK values. *B*, the full dependence of pmTORC1 + pmTORC2 on V_IR and AMPK. The data points corresponding to those shown in panel *A* are indicated by red diamonds (V_IR = 0.00737) and blue squares (V_IR = 0.2).

For a given AMPK activation level, increasing nutrient availability (V_IR) generally promotes cell population growth. That is intuitive and not at all surprising. A more interesting consideration is how AMPK activation may impact tumor cell population at different nutrient levels. Activation of AMPK facilitates the phosphorylation of mTORC2 but inhibits mTORC1. What is then the combined effect of AMPK activation on cancer cell proliferation and survival? Interestingly, the answer depends on the cell microenvironment, specifically, on the nutrient level of the cancer environment. At low nutrient levels, which corresponds to low insulin receptor activation (V_IR = 0.00737 in Fig. 3), increasing AMPK abundance from 20 to 480 increases phosphorylated mTORC2 by 3.87 folds, but has relatively negligible effect on mTORC1 activation, despite AMPK’s general inhibitory effect on mTORC1. This is because the inhibition of mTORC1 by AMPK is partially offset by the indirect activation of mTORC1 by mTORC2. Taken together, enhancing AMPK abundance results in a 13% increase in total phosphorylated mTORC1 and mTORC2, which suggest that AMPK activation may have the detrimental effect on promoting the cancer cell population. See the left set of bars in Fig. 3A and the red diamonds in Fig. 3B. In contrast, in a microenvironment with sufficiently high nutrient levels (V_IR = 0.02), increasing AMPK abundance from 20 to 480 reduces the total phosphorylated mTORC1 and mTORC2 by 22% (right set of bars in Fig. 3A and the blue squares in Fig. 3B). Thus, for cancer cells with chronic nutrient deprivation, AMPK inactivation is expected to have the beneficial effect of limiting their population.

### SIRT1 inhibition produces markedly different cellular dynamics depending on cellular nutrient level

Model results above suggest that under certain cancer conditions, AMPK suppression may limit cancer cell population. This result is consistent with the observation that AMPK activation promotes prostate cancer cell growth under glucose deprivation (31). However, while AMPK activators such as metformin are widely available, the same cannot be said for AMPK inhibitors. The AMPK inhibitor Compound C, which has been shown to suppress the growth of some tumors, is limited to laboratory applications (32). Hence, for potential clinical applications we consider an alternative, indirect inhibition of AMPK. SIRT1, a member of the sirtuin protein family, is known as one of the activators of AMPK through the deacetylation of LKB1 (33). SIRT1 can be inhibited by EX-527, which results in mTOR activation and may be beneficial in certain inflammatory injuries (34). Thus, we assessed the effect of SIRT1 inhibition on mTORC1 and mTORC2 dynamics under different cell microenvironments. We simulated the application of a SIRT1 inhibitor at a dosage that reduces SIRT1 bioavailability to 10% of its baseline. We then examined the resulting AMPK and mTORC dynamics under different nutrient levels.

The predicted time-courses for phosphorylated AMPK, mTORC1, and mTORC2, together with the sum of the latter two, are shown in Fig. 4. These profiles were obtained for three different nutrient levels, at V_IR = 0.1, 0.005, and 0.002687. Each profile was normalized by its peak value in that case. In all three cases, SIRT1 inhibition suppresses AMPK. By 62% under conditions with sufficient nutrients (V_IR = 0.1; Fig. 4A1), and to lesser but still significant extent at lower nutritional levels (52 and 43% at V_IR = 0.005 and 0.002687, respectively; Figs. 4B1 and 4C1). At V_IR = 0.1 (i.e., sufficient nutrients), the decrease in activation of AMPK that follows SIRT1 inhibition enhances mTORC1 activation by 30% and reduces mTORC2 activation by 71% (Figs. 4A2 and 4A3). However, these effects are substantially attenuated at lower nutritional levels, with only 3 and 4% changes in mTORC1 and mTORC2 activation levels, respectively, at V_IR = 0.005 (Figs. 4B2 and 4B3), and with those effects becoming negligible at V_IR = 0.002687 (Figs. 4C2 and 4C3). These results suggest that SIRT1 inhibition may promote cell proliferation and limit survival, but that effect may be insignificant under severe nutrient deprivation. This result may seem inconsistent with model prediction discussed earlier (Fig. 3A), where under nutritional stress, inhibiting AMPK significantly reduces total phosphorylated mTORC1 and mTORC2. That discrepancy can be explained, in large part, by the magnitude of AMPK reduction: in Fig. 3A, AMPK activation decreases by 24 folds, which may be achievable by an inhibitor that directly targets AMPK; whereas in Fig. 4C, indirect inhibition of AMPK only reduces its activation by 43%.

**Figure 4.**
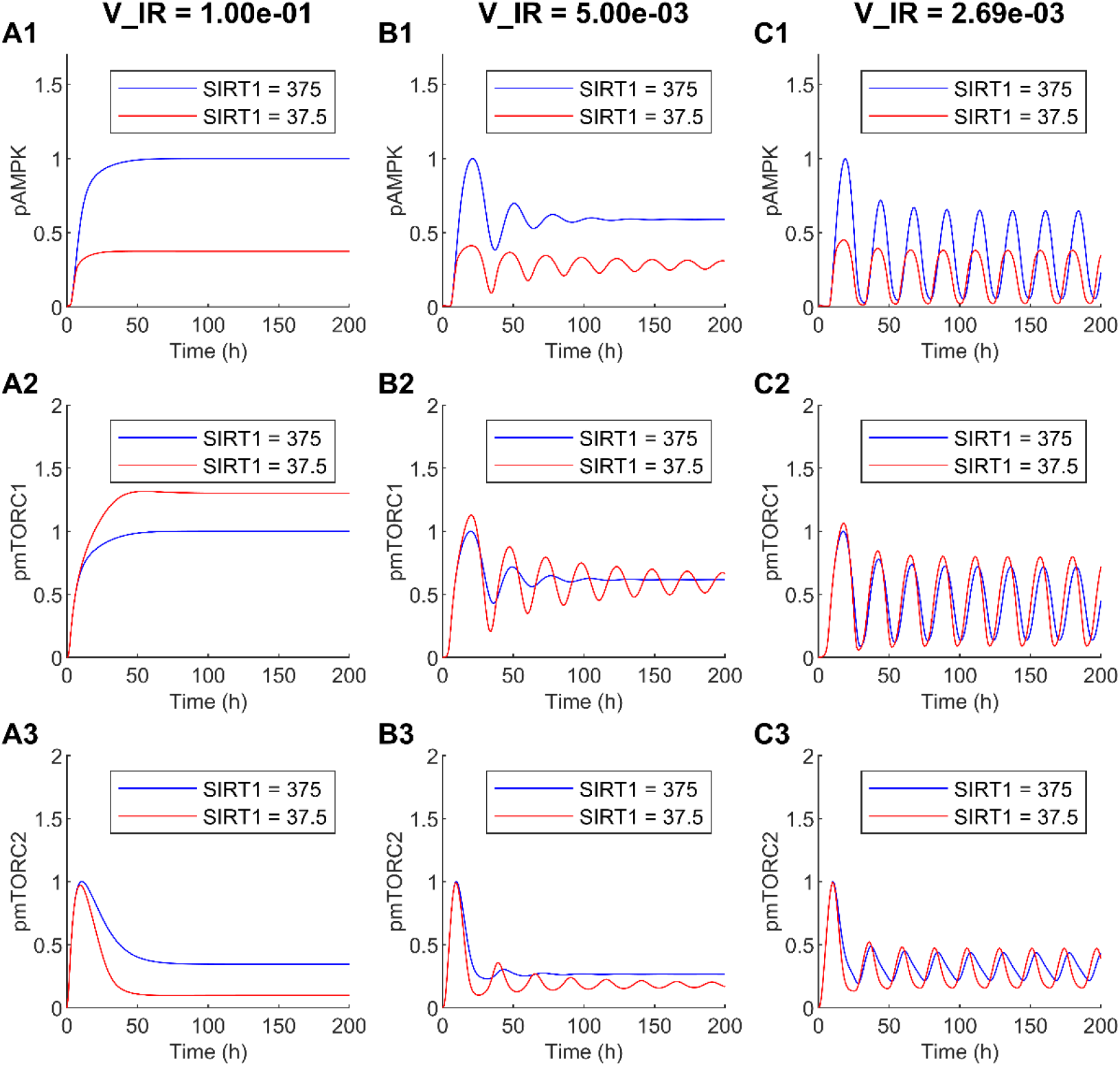
Effect of SIRT1 inhibition on AMPK and mTORC dynamics. *A1--A3*, V_IR = 0.1, which corresponds to high nutrient levels, the model predicts a time-independent steady-state solution. *B1--B3*, V_IR = 0.005, damped oscillations. *C1--C3*, V_IR = 0.00269, sustained oscillations.

Another noteworthy finding is that both V_IR and SIRT1 are bifurcation parameters. Under nutrient deprivation (V_IR = 0.002687), the model predicts sustained oscillations in the protein levels (Fig. 4C), driven by the multitude of feedback loops in the signalling network. Those oscillations may persist, albeit transiently, with higher nutrient availability if SIRT1 is inhibited (V_IR = 0.005, Fig. 4B). At sufficiently high nutritional levels, the model predicts time-independent steady-state solutions (V_IR = 0.1, Fig. 4A).

### Effect of concomitant changes in DEPTOR and AMPK levels

DEPTOR is another key regulator of the mTORC family. We seek to understand how changes in AMPK and DEPTOR abundance shift the balance between cell proliferation and survival, measured by the mTORC1/mTORC2 ratio, and how that shift is affected by cellular environments. With the plentitude of feedback loops and other connections among the signalling proteins, the model predicts that variations in AMPK and DEPTOR abundance may generate complex mTORC dynamics. Results, obtained for V_IR = 0.01 and 0.002687, are shown in Fig. 5.

**Figure 5.**
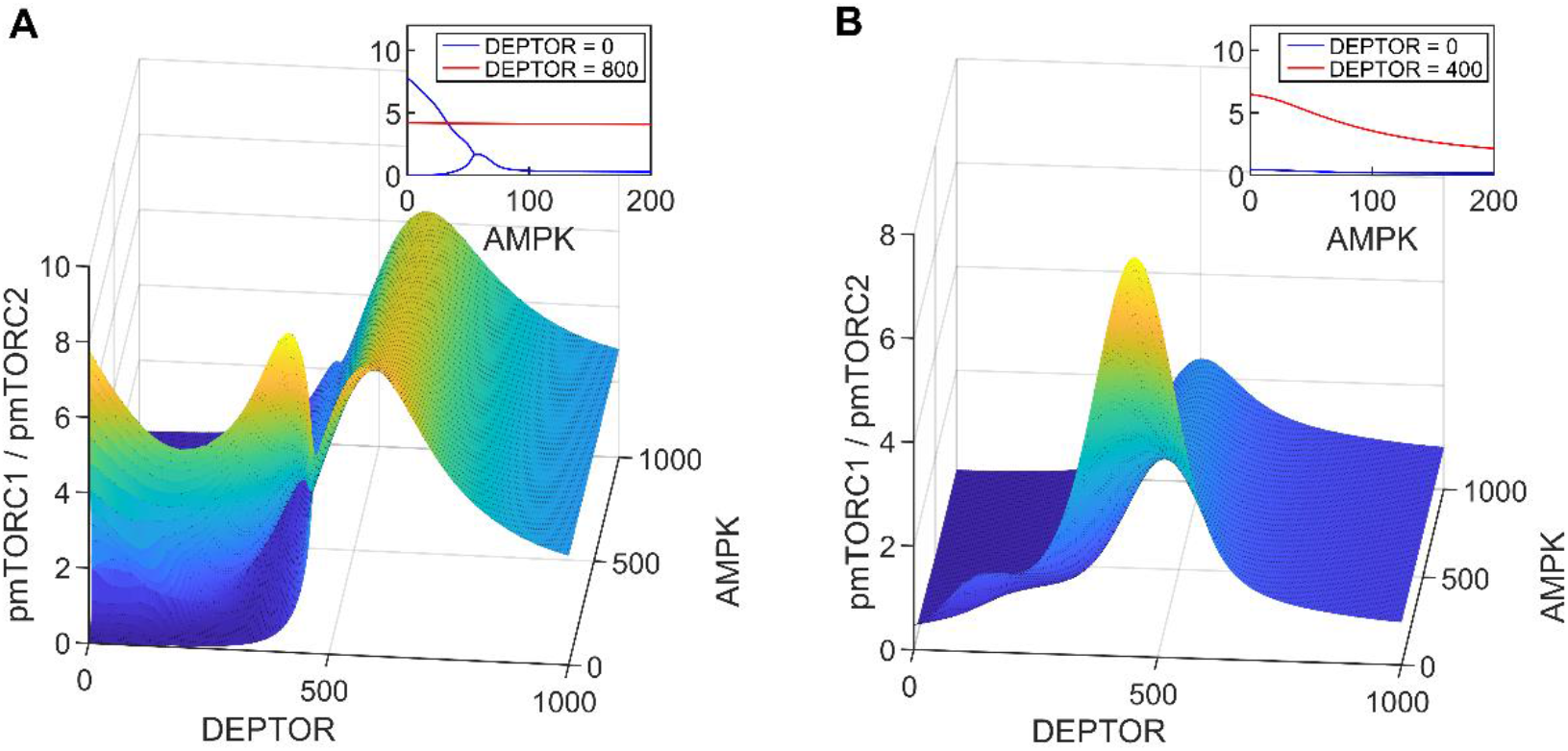
The shift between cell proliferation (pmTORC1) and cell survival (pmTORC2) as one moves around the DEPTOR and AMPK parameter space. *A*, V_IR = 0.002687. *B*, V_IR = 0.01. Insets show ratios of pmTORC1/pmTORC2 as functions of AMPK, at specific DEPTOR values. Inset in Fig. 5A shows that oscillations are predicted for V_IR = 0.002687, DEPTOR = 0, and low AMPK values.

DEPTOR inhibits both mTORC1 and mTORC2 but to different degrees, and the feedback loops introduce further complexity. With sufficient nutrients and in the absence of stress in the cellular environment (V_IR = 0.01, Fig. 5B), the model predicts that cell proliferation is maximized at mid-range DEPTOR abundance values (~400-500, see Fig. 5B); either decreasing or increasing DEPTOR abundance favors cell survival over proliferation, as does increasing AMPK abundance (see Fig. 5B inset). mTORC dynamics is most sensitive to variations in AMPK abundance at this mid-range DEPTOR abundance regime; at sufficiently low or high DEPTOR abundance, changing AMPK abundance does not noticeably shift the cell proliferation versus survival balance. Sustained oscillations are not observed in the parameter space considered.

Under nutrient deprivation or stress (V_IR = 0.002687, Fig. 5A), sustained oscillations are predicted at sufficiently low DEPTOR and AMPK abundance, but not in other parameter regimes. With low DEPTOR abundance, increasing AMPK abundance introduces a significant shift towards cell survival (Fig. 5A inset). However, as DEPTOR abundance increases, AMPK’s role in the competition between cell proliferation and survival becomes negligible, as can be seen in the red line in Fig. 5A inset, which shows that the pmTORC1/pmTORC2 ratio remains essentially unchanged as AMPK increases from 0 to 200.

### Simultaneous suppression of cell proliferation and survival in cancer

To discourage cancer cell survival, mTORC2 may be inhibited via pharmaceutical manipulations. However, given the many protein-protein interactions, inhibition of mTORC2 inevitably affects mTORC1 activity, with potential unfavorable consequences. To understand the synergistic effect on cell proliferation and survival, we simulated the effect of an mTORC2 inhibitor, by varying the rate of deactivation of mTORC2 (V_pmTORC2). Given the somewhat unintuitive effect of AMPK activation on cell growth identified above under nutrient deprivation, model simulations were conducted for a range of AMPK abundance with V_IR = 0.002687. In this parameter regime, model solutions are oscillatory. The two surfaces shown in Fig. 6 correspond to the maxima and minima of the solutions. Additional response curves for mTORC1 and mTORC2 are shown in Figs. 3S and 4S.

**Figure 6.**
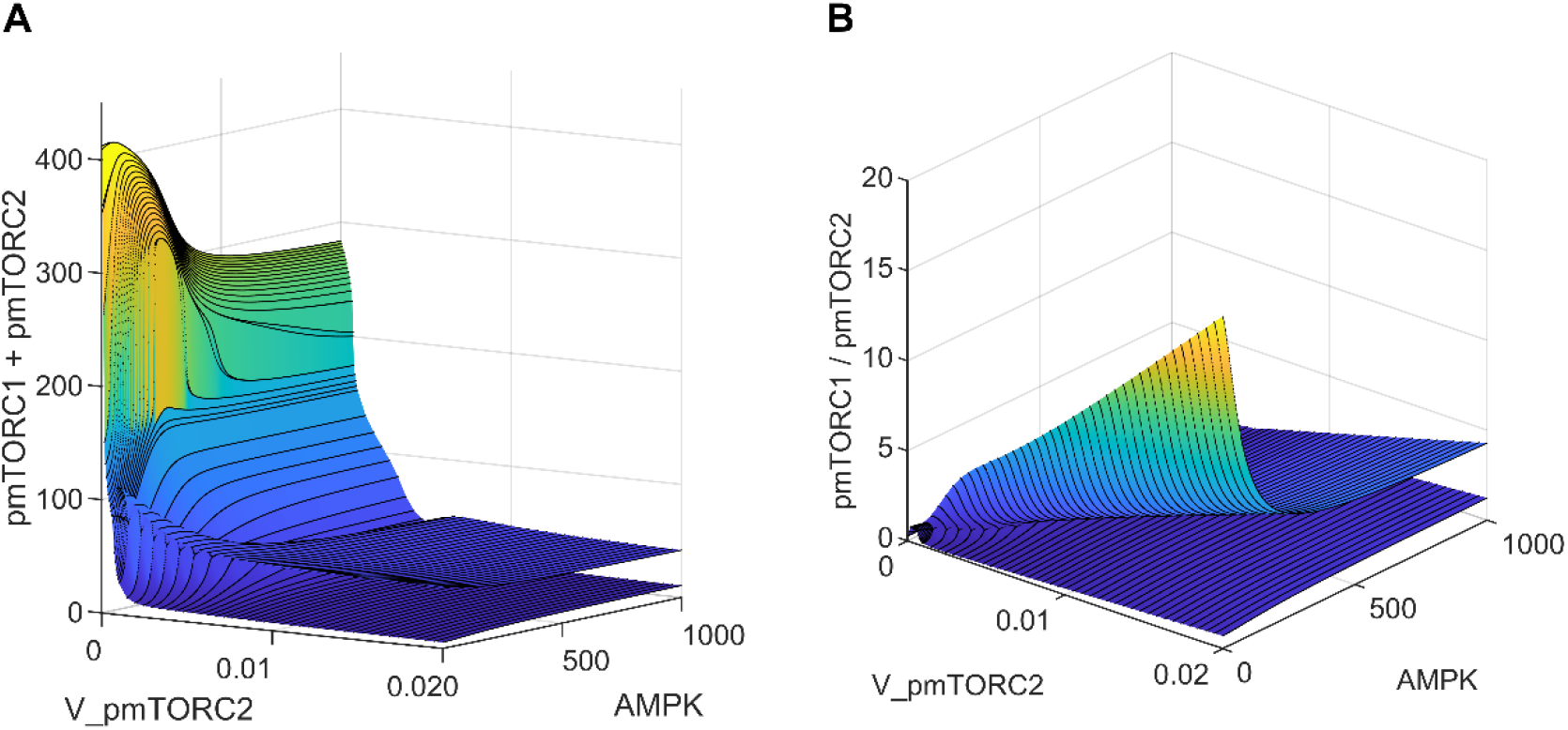
The effect of mTORC2 inhibition and AMPK abundance on cell proliferation and survival. The two surfaces represent the envelope of an oscillatory quantity.

Inhibition of mTORC2, which corresponds to high V_pmTORC2 values, inhibits mTORC1 via AKT deactivation. Under nutritional deprivation, the competing effects on the two mTOR complexes yield an overall reduction in total phosphorylation of mTORC1 and mTORC2; see Fig. 6A. This trend is observed for the range of AMPK abundance considered. However, that comes with the potentially detrimental effect of enhancing cell proliferation, as evinced by the increasing pmTORC1/pmTORC2 ratio as V_pmTORC2 increases (Fig. 6B) for a fixed AMPK. As such, to simultaneously suppress both cell proliferation (mTORC1) and survival (mTORC2), one may combine an mTORC2 inhibitor with an AMPK activator. At sufficiently high V_pmTORC2 and AMPK values, both the sum and ratio of phosphorylated mTORC1 and mTORC2 attain their minima (Fig. 6B).

## Discussion

As a cellular energy sensor, AMPK is activated in response to conditions that deplete cellular energy levels, such as nutrient starvation (especially glucose), hypoxia, and exposure to toxins that inhibit the mitochondrial respiratory chain complex (35). Thus, AMPK plays a key role in coordinating metabolic pathways and in balancing nutrient supply with energy demand. Because of the favorable physiological outcomes of AMPK activation on metabolism, AMPK is believed to have therapeutic importance for treating obesity, type 2 diabetes, non-alcoholic fatty liver disease, and cardiovascular disease (36).

A number of roles have been hypothesized for AMPK in tumorigenesis, both as a promoter and a suppressor. A potential role for AMPK in limiting tumorigenesis is supported by its activation at low ATP by LKB1 (18), which is a tumor suppressor gene that is mutationally inactivated in a number of cancers. Administration of AMPK activators has demonstrated anti-tumorigenesis effects in culture and mice, and in some genetic contexts (37–39). Activated AMPK phosphorylates downstream targets that activate catabolic pathways, while switching off anabolic pathways and other ATP-consuming processes. As a result, AMPK not only promotes ATP synthesis but also restricts cell growth and proliferation in an attempt to restore energy homeostasis and maintain cell viability. However, a conflicting role for AMPK emerges as it is found to promote cell survival under nutrient-poor conditions. This effect is particularly relevant for cancer cells, which are often challenged with insufficient nutrients in the microenvironments to support their needs. Thus, AMPK is also hypothesized to play a pro-tumorigenic role and its presence may be essential to sustain the rapid growth of some cancer populations. Indeed, findings in cancer cell lines and orthotopic xenografts (25) suggest that AMPK is required for some tumor cells to survive under metabolic stress (31,40).

Is AMPK beneficial or malevolent in cancer? More specifically, does AMPK promote or limit cancer cell proliferation and survival? This question is challenging to answer due to the complexity of the signalling pathways that regulate cell growth, which involve many positive and negative feedback loops. To attain insights into the synergy among these processes, and to unravel the effect of activating or inhibiting AMPK under different cellular microenvironments, we have developed the present computational model. To interrogate the effect of AMPK activation on cancer cell population, we applied the model to assess the effect on mTORC1, the activation of which promotes cell proliferation, and mTORC2, the activation of which favors cell survival (26). Model simulations (Fig. 3) suggest that under nutrient-poor conditions, AMPK activation may have an overall pro-tumorigenic role by facilitating cellular survival and hence the growth of cancer. In contrast, with sufficient nutrient availability, the anti-proliferation effect of AMPK dominates, and AMPK acts as a tumor suppressor. Taken together, whether AMPK is pro- or anti-tumorigenic depends, in part, on the nutrient level of the microenvironments of the cancer cells.

Model prediction is consistent with findings in a study by Saito et al. (41), which investigated the extent to which AMPK is critical in achieving metabolic homeostasis in leukemia-initiating cells. They observed that AMPK deletion caused a drastic loss in leukemia cells in the bone marrow, a nutrient-poor environment, but that effect is substantially attenuated in leukemia cells in the spleen, where nutrients are relatively plentiful (41). These findings have potentially groundbreaking implications in cancer therapies. Before the onset of cancer, there was clearly no concern regarding the effect of AMPK on cancer cell survival. Hence, the health impacts of AMPK activators such as metformin are clearly positive. But in established cancers, AMPK can be a double-edged sword (42). In most cancers, the affected organs (e.g., spleen) are well perfused and cancer cells have access to sufficient nutrients; under these conditions, AMPK opposes cancer growth and proliferation. But in some cancers, the microenvironments are nutrient-poor (e.g., bone marrow leukemia), and cancer cells are more dependent on AMPK activity. In that case, the activation of AMPK would increase the viability of the tumour cells and thereby potentially decrease survival of the patient. Thus, in cancers such as bone marrow leukemia, it would be an AMPK inhibitor rather than an activator that might be therapeutically useful.

Because AMPK inhibitors are not widely available, we conducted simulations to explore alternative approaches to reduce cancer cell population. We considered indirect AMPK inhibition via the inhibition of SIRT1, which is required for AMPK activation (22). However, while SIRT1 inhibition influences mTORC1 and mTORC2 activation levels, those effects are insignificant when nutrients are scarce (Fig. 4). Given that this is the cell microenvironment in which AMPK inhibition may limit cancer cell population growth (Fig. 3), what is an alternative means of simultaneously suppressing mTORC1 and mTORC2, if not via SIRT1 inhibition? The predicted negligible effect of SIRT1 inhibition may be attributable to the significant but insufficiently large impact on AMPK activation. With a 90% reduction in SIRT1 abundance, the model predicts less than 50% reduction in AMPK activation under nutrient deprivation, due to the opposing effect from the mTORC1-ULK1-AMPK feedback loop (see Fig. 7). In contrast, when AMPK is much more strongly inhibited, the model predicts significant changes in mTORC1 and mTORC2 (Fig. 3A). An implication of these results is that, if one seeks to combat a cancer whose growth is known to be promoted by AMPK activation, indirect inhibition of AMPK may not be sufficient due to the many compensatory feedback loops in the network. Direct AMPK inhibition may be required. Compound C is available as an AMPK inhibitor and has been demonstrated as a tumor suppressor for certain cancer types (43). However, due to its toxicity Compound C is currently limited to laboratory applications. Hence, cancer research may benefit from enhanced effort in the development of a clinically usable AMPK inhibitor.

**Figure 7.**
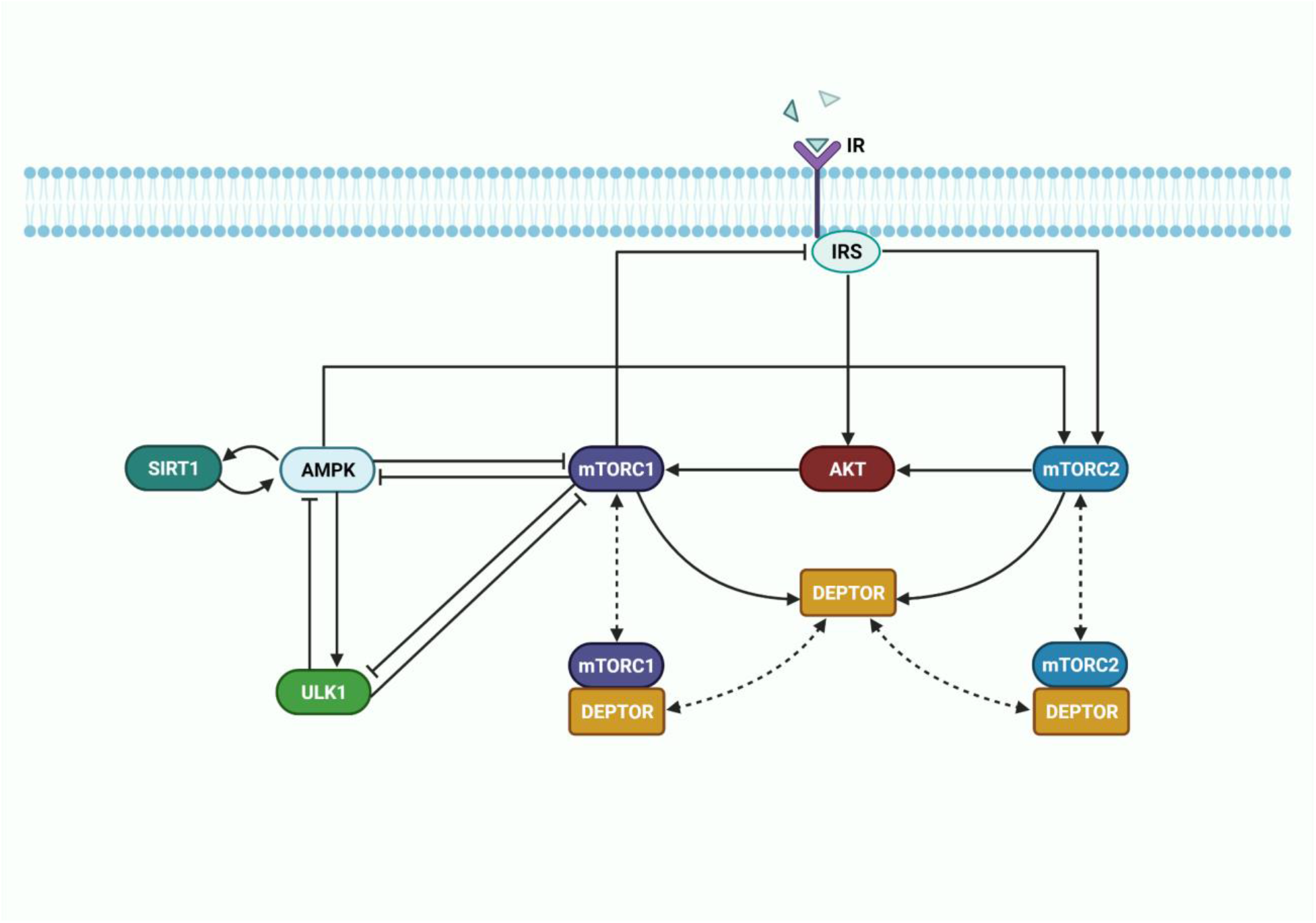
Model metabolic signalling network. A schematic diagram depicting the interactions and feedback loops within the DEPTOR-mTOR network and their connections with external inhibitors and activators AMPK, ULK1, and insulin receptor substrate (IRS). Normal, blunt and dashed arrows denote activation, inhibition, and complex formation, respectively.

The complexity of AMPK’s role as a target in anti-cancer therapy is due, in large part, to its activation of mTORC2, which enhances cell survival (26). This fact points to the potential of mTORC2 inhibitors in cancer treatment. Indeed, selective mTORC2 inhibition has shown promise in blocking breast cancer cell growth and survival and in slowing the migration and metastasis of melanoma cells (44). However, given the many protein-protein interactions, inhibition of mTORC2 inevitably affects mTORC1 activity, possibly with detrimental outcomes. Furthermore, excessive inhibition of mTORC2 may overly suppress AKT, which is essential in the translocation of GLUT4, and lead to insulin insensitivity (45). Our simulations indicate that if an mTORC2 inhibitor is used in an anti-cancer therapy, its effect in combating the progression of cancer will be enhanced by combining with an AMPK activator (Fig. 6).

Another notable protein that modulates mTOR signalling is DEPTOR, which associates with both mTORC1 and mTORC2 and physically interacts with the FAT domain of mTOR through its PDZ domain (46). The role of DEPTOR as a tumor suppressor is consistent with its low expression revealed in a number of human cancers, including pancreas (12), esophageal squamous cell carcinoma (13), lungs (16), and breast cancer (14). The role of DEPTOR in inhibiting tumor progression may be attributed to its ability to repress cell migration, which is necessary in metastasis and cancer progression. Supportive evidence is provided by the significantly reduced DEPTOR expression reported on the invasive front of endometrial cancer tissues. Consistency with the multifaceted nature of most aspects of cancer, DEPTOR has been reported to be overexpressed in multiple myeloma cells (10), which suggests a role of oncogene rather than tumor suppressor. Whether and how DEPTOR might promote metastasis in specific contexts awaits clarification in future studies. Our model simulations suggest that the relative effect of DEPTOR on cancer cell proliferation and survival depends on its expression level and other factors, including metabolic stress and AMPK activation level (Fig. 5).

Given the complexity of the signalling network, the effect of a single perturbation is often difficult to second guess. The cascading effects and feedback response can be fully explored using the present model. For example, the DEPTOR simulations suggest that with adequate nutrients, administering either a DEPTOR activator or inhibitor can shift cell fate away from proliferation in favor of cell survival; but in nutrient deprivation, DEPTOR inhibition may introduce oscillations in mTORC (Fig. 5). Another potentially impactful application of the model lies in the exploration of drug-drug interactions. In the mTORC2 inhibitor and AMPK activator simulations, the model predicts that when co-administered, these drugs may effectively limit cancer population and the effect of both drugs on the system is not additive and is difficult to predict without a mathematical model (Fig. 6).

### Limitations of study

A major limitation of the present study is that while the model reproduces cell culture experimental results, its predictions may well deviate from *in vivo* observations. Also, molecular alterations involving the mTOR pathway have been reported in a number of cancer types (47), which may limit the implications of the model’s predictions to cancers. Despite its limitations, with appropriate extensions the model may yet prove useful in investigations of cell growth and metabolism in mammals, in both health and disease. Indeed, the model is applicable to many diseases other than cancer (48). One example is diabetes. One novel anti-hyperglycemic medication is the sodium-glucose cotransporter 2 (SGLT2) inhibitor, which targets the kidney (49). SGLT2 inhibition in diabetes is known to delay the progression toward or of diabetic kidney disease and offer cardiovascular protection (50). An additional benefit demonstrated at least in the SGLT2 inhibitor canagliflozin is its activation of AMPK (51), which may have positive effects on kidney metabolism. To study how the administration of canagliflozin affects renal hemodynamics and bioenergetics in a patient with diabetes, one may incorporate the present model into a computational model of kidney function and metabolism (52–54).

The current set of parameters are fitted for diverse physiological settings. Model parameters can be modified to study a specific cell type and condition. One important variable is sex (55). For instance, sex-specific models can be formulated to investigate the functional implications of the differential mTORC1 and mTORC2 activity levels reported in male and female mice (56). Age is another worthwhile direction. Depending on the age of the individuals, mTORC1 in skeletal muscle cells is activated under different conditions (57,58). The development of models that take into account cell or tissue specificity, together with sex and age, can help identify therapeutic strategies that target metabolic pathways, for individualized treatments.

## Materials and Methods

The model represents key proteins in cellular metabolism: insulin receptor substrate (IRS) and AKT, which modulate most of insulin’s effects on metabolism; mTORC1, mTORC2, DEPTOR, and ULK1, which mediate mTORC1’s signal in autophagy initiation; and the two key energy sensors, AMPK and SIRT1. The dynamics of the signaling pathways is modeled as a system of ordinary differential equations that involve Michaelis Menten and mass action kinetics. A schematic diagram of the model signalling pathway is depicted in Fig. 7. Model reactions are presented in Table 1S (supplemental materials).

Insulin activates the insulin receptor (IR), which triggers the IRS, resulting in the phosphorylation of mTORC2. mTORC2 phosphorylates AKT (48). Activation of IRS also independently activates AKT, which activates mTORC1 through the phosphorylation of the tumor suppressor TSC2 (not represented). Activation of mTORC1 has a number of downstream effects, including a negative feedback loop to IRS, and the inhibition of ULK1 (3,8). The ULK1 complex is a key contributor to the initiation of autophagy.

Both mTORC1 and mTORC2 are modulated by AMPK, a master regulator of cellular energy that is activated under starvation or hypoxia. AMPK can be stimulated by LKB1 (not represented), which is deacetylated by SIRT1. Activated AMPK promotes autophagy by directly phosphorylating and activating ULK1 (59). As such, there is a competition between mTORC1 and AMPK to phosphorylate different residues of ULK1 to decide cell fate. ULK1 in turn inhibits AMPK and mTORC1 in a negative feedback loop, whereas leucine activates them. These feedback loops and interactions are summarized in Fig. 7.

### Parameter Estimation

Given that our main objective is to characterize possible emergent properties of the target metabolic network under diverse physiological settings, we aim to explore the network behaviour over wide ranges of kinetic parameters, rather than constraining them to a specific dataset from a particular experimental model (60). Nonetheless, model parameters are constrained by biologically plausible values. To determine the baseline parameter set, we perform careful calibration using data from multiple experimental studies. Specifically, model parameters are calibrated to satisfy the following criteria simultaneously: (i) the model predicts oscillations in mTORC1, ULK1 and AMPK under energetic stress that approximate experimental data in Ref. (27), (ii) the feedback response of mTORC1 to variations in AMPK approximate experimental data in Ref. (28), and (iii) the model predicts changes in AMPK and AKT activation in response to treatment of AICAR, an AMPK activator, consistent with experimental data in Ref. (26,29). We seek to satisfy criteria (i)--(iii) simultaneously via an iterative process. Criterion (i) is satisfied by optimizing the predicted time-profiles using the interior point algorithm in the MATLAB built-in function fmincon. Criterion (ii) is satisfied by adjusting parameters K_AMPK and K_AMPK_by_SIRT1 to minimize the least-square difference between the predicted mTORC1 and mTORC2 response curves, as the fractional activation of AMPK varies from 0.1 to 1.0. Criterion (iii) is satisfied by adjusting parameters K_AMPK and K_AMPK_by_SIRT1 to minimize the difference between the measured and predicted AMPK and AKT activation levels. These procedures are repeated iteratively until all three criteria are satisfied simultaneously. Baseline model parameters are given in Table 2S (supplemental materials).

### Bifurcation Analysis

To assess the impact of insulin receptor activation, [and whatever V_pmTORC2 and K_mTORC2_DEPTOR_diss describe], and AMPK abundance on metabolism, we conducted a bifurcation analysis to assess model sensitivity to variations in parameters V_IR, V_pmTORC2, K_mTORC2_DEPTOR_diss, and AMPK abundance.The parameters are varied and at each point, the system of ODEs is solved in MATLAB from t = 0 to 1000 hours. The timespan is set to last up to 1000 hours to ensure that the system has reached steady state or limit cycle oscillation. The maximum and minimum protein concentrations of the phosphorylated forms of mTORC1 and mTORC2 are determined from the last 200 hours in time. Subsequently, they are plotted as single points in the bifurcation diagram. Plotting both the maximum and the minimum allows the identification of limit cycle oscillation, as this would capture the maxima and minima of oscillation.

To understand the interplay between cell environment and type (shows itself as insulin sensitivity of the cell) and the amount of AMPK both V_IR and AMPK abundance were changed simultaneously. As a result, three-dimensional bifurcation diagrams are plotted. A 1000 by 1000 mesh is defined with varying V_IR and AMPK abundance, and the initial condition for AMPK abundance varying from 0 to 1000. At each point in the mesh, the maximum and minimum protein concentrations are plotted following the same procedure described previously.

### Local sensitivity analysis

Local sensitivity analysis is conducted on mathematical models to assess how small perturbations in individual model parameters affect the output signals. The local sensitivity of a steady-state output signal S to changes in the parameter p can be determined by computing the normalized derivative of S with respect to p:

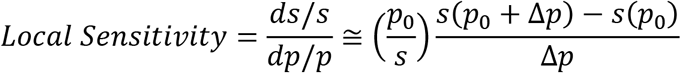

S(p_0_) corresponds to the steady state concentration of the signal S obtained using the initial parameter set p_0_. The parameter of interest p is perturbed by 0.5 % (i.e., Δp = 0.005 * p_0_), and the new steady state concentration S(p_0_+Δp) is obtained.

### Global sensitivity analysis

In the global sensitivity analysis, the response output variables to perturbations in parameter values is investigated across the entire parameter space. Model parameters are varied over the physiological ranges, where the maximum velocity (V_max_) ranges from 0.001 to 10 nM/s, rates of formation and dissociation (K_form, K_diss) range from 10^−6^ to 10 nM^−1^s^−1^, rate of catalytic reaction (K) ranges from 10^−6^ to 1 s^−1^, and the Michaelis-Menten constant (Km) ranges from 1 to 1000 nM. N = 100000 parameter sets are randomly sampled from the parameter space using Latin hypercube sampling.

Model equations are solved for each parameter set p_0_ to yield steady-state solution S. The partial rank correlation coefficient (PRCC) is computed to characterize the correlation between p_0_ and S using the following equation:

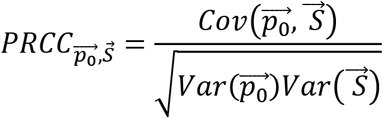

## Data availability

The datasets and computer code produced in this study are available in https://github.com/MehrshadSD/

## Acknowledgements

This study is partially supported by the Canada 150 Research Chairs Program and the Natural Sciences and Engineering Research Council of Canada (NSERC Discovery award: RGPIN-2019-03916);

## Author contributions

Conceptualization, MS; methodology, MS and DS; software, DS; validation, MS and DS; formal analysis, MS, DS and ATL; resources, ATL; data curation, MS, DS; writing— original draft preparation, MS and ATL; writing—review and editing, MS and ATL; visualization, DS; supervision, MS and ATL; project administration, ATL; funding acquisition, ATL. All authors have read and agreed to the published version of the manuscript.

## Conflict of interest

The authors declare no conflict of interest.

## Bibliography

1. DeBerardinis RJ, Chandel NS. Fundamentals of cancer metabolism. Sci Adv. 2016 May 27;2(5):e1600200.

2. Ward PS, Thompson CB. Metabolic reprogramming: a cancer hallmark even warburg did not anticipate. Cancer Cell. 2012 Mar 20;21(3):297–308.

3. Saxton RA, Sabatini DM. mTOR Signaling in Growth, Metabolism, and Disease. Cell. 2017 Mar 9;168(6):960–76.

4. Forbes SA, Bindal N, Bamford S, Cole C, Kok CY, Beare D, et al. COSMIC: mining complete cancer genomes in the Catalogue of Somatic Mutations in Cancer. Nucleic Acids Res. 2011 Jan;39(Database issue):D945–50.

5. Hanahan D, Weinberg RA. Hallmarks of cancer: the next generation. Cell. 2011 Mar 4;144(5):646–74.

6. Pópulo H, Lopes JM, Soares P. The mTOR signalling pathway in human cancer. Int J Mol Sci. 2012 Feb 10;13(2):1886–918.

7. Benjamin D, Colombi M, Moroni C, Hall MN. Rapamycin passes the torch: a new generation of mTOR inhibitors. Nat Rev Drug Discov. 2011 Oct 31;10(11):868–80.

8. Sadria M, Layton AT. Interactions among mTORC, AMPK and SIRT: a computational model for cell energy balance and metabolism. Cell Commun Signal. 2021 May 20;19(1):57.

9. Gruppuso PA, Boylan JM, Sanders JA. The physiology and pathophysiology of rapamycin resistance: implications for cancer. Cell Cycle. 2011 Apr 1;10(7):1050–8.

10. Peterson TR, Laplante M, Thoreen CC, Sancak Y, Kang SA, Kuehl WM, et al. DEPTOR is an mTOR inhibitor frequently overexpressed in multiple myeloma cells and required for their survival. Cell. 2009 May 29;137(5):873–86.

11. Varusai TM, Nguyen LK. Dynamic modelling of the mTOR signalling network reveals complex emergent behaviours conferred by DEPTOR. Sci Rep. 2018 Jan 12;8(1):643.

12. Li H, Sun GY, Zhao Y, Thomas D, Greenson JK, Zalupski MM, et al. DEPTOR has growth suppression activity against pancreatic cancer cells. Oncotarget. 2014 Dec 30;5(24):12811–9.

13. Ji Y-M, Zhou X-F, Zhang J, Zheng X, Li S-B, Wei Z-Q, et al. DEPTOR suppresses the progression of esophageal squamous cell carcinoma and predicts poor prognosis. Oncotarget. 2016 Mar 22;7(12):14188–98.

14. Parvani JG, Davuluri G, Wendt MK, Espinosa C, Tian M, Danielpour D, et al. Deptor enhances triple-negative breast cancer metastasis and chemoresistance through coupling to survivin expression. Neoplasia. 2015 Mar;17(3):317–28.

15. Hu B, Lv X, Gao F, Chen S, Wang S, Qing X, et al. Downregulation of DEPTOR inhibits the proliferation, migration, and survival of osteosarcoma through PI3K/Akt/mTOR pathway. Onco Targets Ther. 2017 Sep 8;10:4379–91.

16. Zhou X, Guo J, Ji Y, Pan G, Liu T, Zhu H, et al. Reciprocal Negative Regulation between EGFR and DEPTOR Plays an Important Role in the Progression of Lung Adenocarcinoma. Mol Cancer Res. 2016 Feb 19;14(5):448–57.

17. Vincent EE, Coelho PP, Blagih J, Griss T, Viollet B, Jones RG. Differential effects of AMPK agonists on cell growth and metabolism. Oncogene. 2015 Jul;34(28):3627–39.

18. Hawley SA, Boudeau J, Reid JL, Mustard KJ, Udd L, Mäkelä TP, et al. Complexes between the LKB1 tumor suppressor, STRAD alpha/beta and MO25 alpha/beta are upstream kinases in the AMP-activated protein kinase cascade. J Biol. 2003 Sep 24;2(4):28.

19. Steinberg GR, Carling D. AMP-activated protein kinase: the current landscape for drug development. Nat Rev Drug Discov. 2019;18(7):527–51.

20. Carling D. AMPK signalling in health and disease. Curr Opin Cell Biol. 2017 Feb 21;45:31–7.

21. Steinberg GR, Schertzer JD. AMPK promotes macrophage fatty acid oxidative metabolism to mitigate inflammation: implications for diabetes and cardiovascular disease. Immunol Cell Biol. 2014 Apr;92(4):340–5.

22. Price NL, Gomes AP, Ling AJY, Duarte FV, Martin-Montalvo A, North BJ, et al. SIRT1 is required for AMPK activation and the beneficial effects of resveratrol on mitochondrial function. Cell Metab. 2012 May 2;15(5):675–90.

23. Faubert B, Boily G, Izreig S, Griss T, Samborska B, Dong Z, et al. AMPK is a negative regulator of the Warburg effect and suppresses tumor growth in vivo. Cell Metab. 2013 Jan 8;17(1):113–24.

24. Jeon S-M, Chandel NS, Hay N. AMPK regulates NADPH homeostasis to promote tumour cell survival during energy stress. Nature. 2012 May 9;485(7400):661–5.

25. Kishton RJ, Barnes CE, Nichols AG, Cohen S, Gerriets VA, Siska PJ, et al. AMPK Is Essential to Balance Glycolysis and Mitochondrial Metabolism to Control T-ALL Cell Stress and Survival. Cell Metab. 2016 Apr 12;23(4):649–62.

26. Kazyken D, Magnuson B, Bodur C, Acosta-Jaquez HA, Zhang D, Tong X, et al. AMPK directly activates mTORC2 to promote cell survival during acute energetic stress. Sci Signal. 2019 Jun 11;12(585).

27. Holczer M, Hajdú B, Lőrincz T, Szarka A, Bánhegyi G, Kapuy O. Fine-tuning of AMPK-ULK1-mTORC1 regulatory triangle is crucial for autophagy oscillation. Sci Rep. 2020 Oct 20;10(1):17803.

28. Pal PB, Sonowal H, Shukla K, Srivastava SK, Ramana KV. Aldose reductase regulates hyperglycemia-induced HUVEC death via SIRT1/AMPK-α1/mTOR pathway. J Mol Endocrinol. 2019 Jul 1;63(1):11–25.

29. Habib SL, Yadav A, Kidane D, Weiss RH, Liang S. Novel protective mechanism of reducing renal cell damage in diabetes: Activation AMPK by AICAR increased NRF2/OGG1 proteins and reduced oxidative DNA damage. Cell Cycle. 2016 Nov 16;15(22):3048–59.

30. Ost A, Svensson K, Ruishalme I, Brännmark C, Franck N, Krook H, et al. Attenuated mTOR signaling and enhanced autophagy in adipocytes from obese patients with type 2 diabetes. Mol Med. 2010 Aug;16(7–8):235–46.

31. Chhipa RR, Wu Y, Mohler JL, Ip C. Survival advantage of AMPK activation to androgen-independent prostate cancer cells during energy stress. Cell Signal. 2010 Oct;22(10):1554–61.

32. Yang W-L, Perillo W, Liou D, Marambaud P, Wang P. AMPK inhibitor compound C suppresses cell proliferation by induction of apoptosis and autophagy in human colorectal cancer cells. J Surg Oncol. 2012 Nov;106(6):680–8.

33. Ruderman NB, Xu XJ, Nelson L, Cacicedo JM, Saha AK, Lan F, et al. AMPK and SIRT1: a long-standing partnership? Am J Physiol Endocrinol Metab. 2010 Apr;298(4):E751–60.

34. Gertz M, Fischer F, Nguyen GTT, Lakshminarasimhan M, Schutkowski M, Weyand M, et al. Ex-527 inhibits Sirtuins by exploiting their unique NAD+− dependent deacetylation mechanism. Proc Natl Acad Sci USA. 2013 Jul 23;110(30):E2772–81.

35. Hardie DG, Ross FA, Hawley SA. AMPK: a nutrient and energy sensor that maintains energy homeostasis. Nat Rev Mol Cell Biol. 2012 Mar 22;13(4):251–62.

36. Day EA, Ford RJ, Steinberg GR. AMPK as a therapeutic target for treating metabolic diseases. Trends Endocrinol Metab. 2017 Aug;28(8):545–60.

37. Evans JMM, Donnelly LA, Emslie-Smith AM, Alessi DR, Morris AD. Metformin and reduced risk of cancer in diabetic patients. BMJ. 2005 Jun 4;330(7503):1304–5.

38. Huang X, Wullschleger S, Shpiro N, McGuire VA, Sakamoto K, Woods YL, et al. Important role of the LKB1-AMPK pathway in suppressing tumorigenesis in PTEN-deficient mice. Biochem J. 2008 Jun 1;412(2):211–21.

39. Pineda CT, Ramanathan S, Fon Tacer K, Weon JL, Potts MB, Ou Y-H, et al. Degradation of AMPK by a cancer-specific ubiquitin ligase. Cell. 2015 Feb 12;160(4):715–28.

40. Jeon S-M, Hay N. The dark face of AMPK as an essential tumor promoter. Cell Logist. 2012 Oct 1;2(4):197–202.

41. Saito Y, Chapple RH, Lin A, Kitano A, Nakada D. AMPK Protects Leukemia-Initiating Cells in Myeloid Leukemias from Metabolic Stress in the Bone Marrow. Cell Stem Cell. 2015 Nov 5;17(5):585–96.

42. Russell FM, Hardie DG. AMP-Activated Protein Kinase: Do We Need Activators or Inhibitors to Treat or Prevent Cancer? Int J Mol Sci. 2020 Dec 27;22(1).

43. Liu X, Chhipa RR, Nakano I, Dasgupta B. The AMPK inhibitor compound C is a potent AMPK-independent antiglioma agent. Mol Cancer Ther. 2014 Mar;13(3):596–605.

44. Guenzle J, Akasaka H, Joechle K, Reichardt W, Venkatasamy A, Hoeppner J, et al. Pharmacological Inhibition of mTORC2 Reduces Migration and Metastasis in Melanoma. Int J Mol Sci. 2020 Dec 22;22(1).

45. Houde VP, Brûlé S, Festuccia WT, Blanchard P-G, Bellmann K, Deshaies Y, et al. Chronic rapamycin treatment causes glucose intolerance and hyperlipidemia by upregulating hepatic gluconeogenesis and impairing lipid deposition in adipose tissue. Diabetes. 2010 Jun;59(6):1338–48.

46. Caron A, Briscoe DM, Richard D, Laplante M. DEPTOR at the nexus of cancer, metabolism, and immunity. Physiol Rev. 2018 Jul 1;98(3):1765–803.

47. Zhang Y, Kwok-Shing Ng P, Kucherlapati M, Chen F, Liu Y, Tsang YH, et al. A Pan-Cancer Proteogenomic Atlas of PI3K/AKT/mTOR Pathway Alterations. Cancer Cell. 2017 Jun 12;31(6):820–832.e3.

48. Sadria M, Layton AT. Aging affects circadian clock and metabolism and modulates timing of medication. iScience. 2021 Apr 23;24(4):102245.

49. Michel MC, Mayoux E, Vallon V. A comprehensive review of the pharmacodynamics of the SGLT2 inhibitor empagliflozin in animals and humans. Naunyn Schmiedebergs Arch Pharmacol. 2015 Aug;388(8):801–16.

50. Layton AT, Vallon V. Cardiovascular benefits of SGLT2 inhibition in diabetes and chronic kidney diseases. Acta Physiol (Oxf). 2018 Apr;222(4):e13050.

51. Hawley SA, Ford RJ, Smith BK, Gowans GJ, Mancini SJ, Pitt RD, et al. The na+/glucose cotransporter inhibitor canagliflozin activates AMPK by inhibiting mitochondrial function and increasing cellular AMP levels. Diabetes. 2016 Sep;65(9):2784–94.

52. Layton AT, Layton HE. A computational model of epithelial solute and water transport along a human nephron. PLoS Comput Biol. 2019 Feb 25;15(2):e1006108.

53. Layton AT, Vallon V. SGLT2 inhibition in a kidney with reduced nephron number: modeling and analysis of solute transport and metabolism. Am J Physiol Renal Physiol. 2018 May 1;314(5):F969–84.

54. Hu R, Layton A. A Computational Model of Kidney Function in a Patient with Diabetes. Int J Mol Sci. 2021 May 29;22(11).

55. Layton AT. His and her mathematical models of physiological systems. Math Biosci. 2021 Jun 10;108642.

56. Baar EL, Carbajal KA, Ong IM, Lamming DW. Sex- and tissue-specific changes in mTOR signaling with age in C57BL/6J mice. Aging Cell. 2016 Feb;15(1):155–66.

57. Francaux M, Demeulder B, Naslain D, Fortin R, Lutz O, Caty G, et al. Aging Reduces the Activation of the mTORC1 Pathway after Resistance Exercise and Protein Intake in Human Skeletal Muscle: Potential Role of REDD1 and Impaired Anabolic Sensitivity. Nutrients. 2016 Jan 15;8(1).

58. Fry CS, Drummond MJ, Glynn EL, Dickinson JM, Gundermann DM, Timmerman KL, et al. Aging impairs contraction-induced human skeletal muscle mTORC1 signaling and protein synthesis. Skelet Muscle. 2011 Mar 2;1(1):11.

59. Kim J, Kundu M, Viollet B, Guan K-L. AMPK and mTOR regulate autophagy through direct phosphorylation of Ulk1. Nat Cell Biol. 2011 Feb;13(2):132–41.

60. Kholodenko BN. Cell-signalling dynamics in time and space. Nat Rev Mol Cell Biol. 2006 Mar;7(3):165–76.

